# Complementarity of ecosystem types drives landscape-wide productivity in North America

**DOI:** 10.1101/2025.11.17.688810

**Authors:** Simon Landauer, Florian Altermatt, Reinhard Furrer, Forest Isbell, Eva Spehn, Pascal A. Niklaus

**Affiliations:** Department of Evolutionary Biology and Environmental Studies, University of Zurich, Zurich, 8057, Switzerland; Department of Aquatic Ecology, Eawag: Swiss Federal Institute of Aquatic Science and Technology, Dübendorf, 8600, Switzerland; Department of Mathematical Modeling and Machine Learning, University of Zurich, Zurich, 8057, Switzerland; Department of Ecology, Evolution, and Behavior, University of Minnesota Twin Cities, Saint Paul, 55108, Minnesota, USA; Swiss Biodiversity Forum, SCNAT, Bern, 3001, Switzerland

## Abstract

Landscape mosaics with a greater diversity of ecosystems tend to be more productive, mirroring the well-established relationship between species diversity and productivity observed in plot-scale biodiversity experiments. However, the mechanisms driving this effect at the landscape scale remain unclear. Here, we analyze a 15-year time series of satellite-derived primary productivity across over 50,000 landscape plots that vary in ecosystem composition. Our results demonstrate that more diverse landscapes are more productive and more predictable under environmental stress, especially drought. Using statistical partitioning, we show that these diversity effects are primarily driven by complementarity, with productivity gains that are broadly shared among ecosystem types rather than being dominated by a few. The specific ecosystem types that contributed most to landscape functioning varied regionally, but their role in driving mixture productivity remained unaffected by drought. These findings extend biodiversity theory to the landscape scale, emphasizing the critical role of higher-order diversity in shaping ecosystem function and informing landscape management.

## Introduction

More species-rich plant communities are more productive on average, and the productivity of such communities is temporally more stable^1–4^. This biodiversity-ecosystem functioning (BEF) research has predominantly been conducted in experiments with artificially created communities in which species composition was systematically varied^5,6^. Also, these BEF relationships have mostly been established within single, relatively homogeneous communities, using metrics of biodiversity based on plant species abundances, for example species richness, or diversity indices based on species identities or their relatedness^7^.

Real-world landscapes are composed of a mosaic of different communities, habitats and ecosystem types^8,9^. The basic elements of such mosaics are relatively homogeneous within themselves and can be referred to as land units^10^. Such land units include ecosystem types such as forest or grassland but may also be anthropogenically dominated like croplands and urban areas. In landscapes composed of different land units, an additional form of diversity manifests: the diversity of the land unit types themselves^11^. In the following, we refer to this type of diversity as ’landscape diversity’, and to the units of land harboring this diversity as ’landscapes’ (Box 1). We do so to distinguish these from the much smaller ’plots’ in species-diversity-focused BEF experiments. Because landscape diversity is an emergent form of diversity that does not occur in the isolated and relatively homogeneous plots of BEF experiments, BEF experiments cannot inform on the functional importance of landscape diversity.

Landscape diversity has received some attention in observational studies comparing plant communities of different species richness^12,13^n these studies, landscape diversity has mostly been treated as environmental context, and the analyses of diversity-functioning relationships have predominantly focused on effects of local species diversity. Using networks of experimental model systems, at least one study has shown that abiotic fluxes between ecosystems can support the functioning of the entire ecosystem network in ways similar to how interspecific interactions in multi-species communities support the functioning of these communities^14^. However, it remains unclear whether such interactions between different ecosystem types indeed promote novel types of diversity effects.

In two recent studies across Switzerland and North America, 17- and 20-year time series of satellite-sensed productivity were combined with land-cover (LC) information, finding that landscapes composed of multiple LC types were, on average, more productive than single-LC landscapes^15,16^. These effects align, phenomenologically, with the relationship between plant species richness and community productivity observed in plot-scale BEF experiments and extend them to land cover diversity and landscape productivity. In both studies, local species diversity data, to the extent that it was available, did not explain the changes in landscape productivity that were observed, suggesting that landscape diversity effects might have been promoted by other mechanisms. Yet, the individual productivity contributions of the different LC types present in mixed LC landscapes could not be assessed, as the satellite remote-sensing product employed (vegetation indices collected using the MODIS satellite instrument) had a spatial resolution (250 m) that was insufficient for this purpose.

Here, we analyzed LC-specific productivity changes that underpin landscape diversity effects, and to this end established a new network of ∼50,000 study landscapes, each 25 ha in size, spanning most of North America between 15° and 69° northern latitude. As metric of landscape functioning, we used 30 m spatial resolution indices of vegetation productivity calculated from Landsat satellite data spanning the years 2008-2022. As metric of landscape diversity, we used the Commission for Environmental Cooperation’s North American land monitoring system map^17^ from 2015 (CEC map, 30-m spatial resolution) and determined LC-type richness (LCR), defined as the number of different LC types present in each landscape. We aggregated the original 19 LC classes of the CEC map into six broad, ecologically distinct LC classes: agriculture, forest, grassland, shrubland, wetland, and urban areas. Following design principles from experimental BEF research^18^, we divided the study area into 290 spatial blocks defined by a combination of a 3° latitude ⨯ 6° longitude grid and of an environmental stratification into biogeographic regions^19^, and systematically established replicate LCR gradients within each of these blocks, similar to how species richness gradients are replicated within the blocks of plot-scale BEF experiments (Fig. 1, Methods). Overall, our study comprised of 49 unique LC-type combinations, with LCR ranging from 1 to 4.

**Fig. 1.**
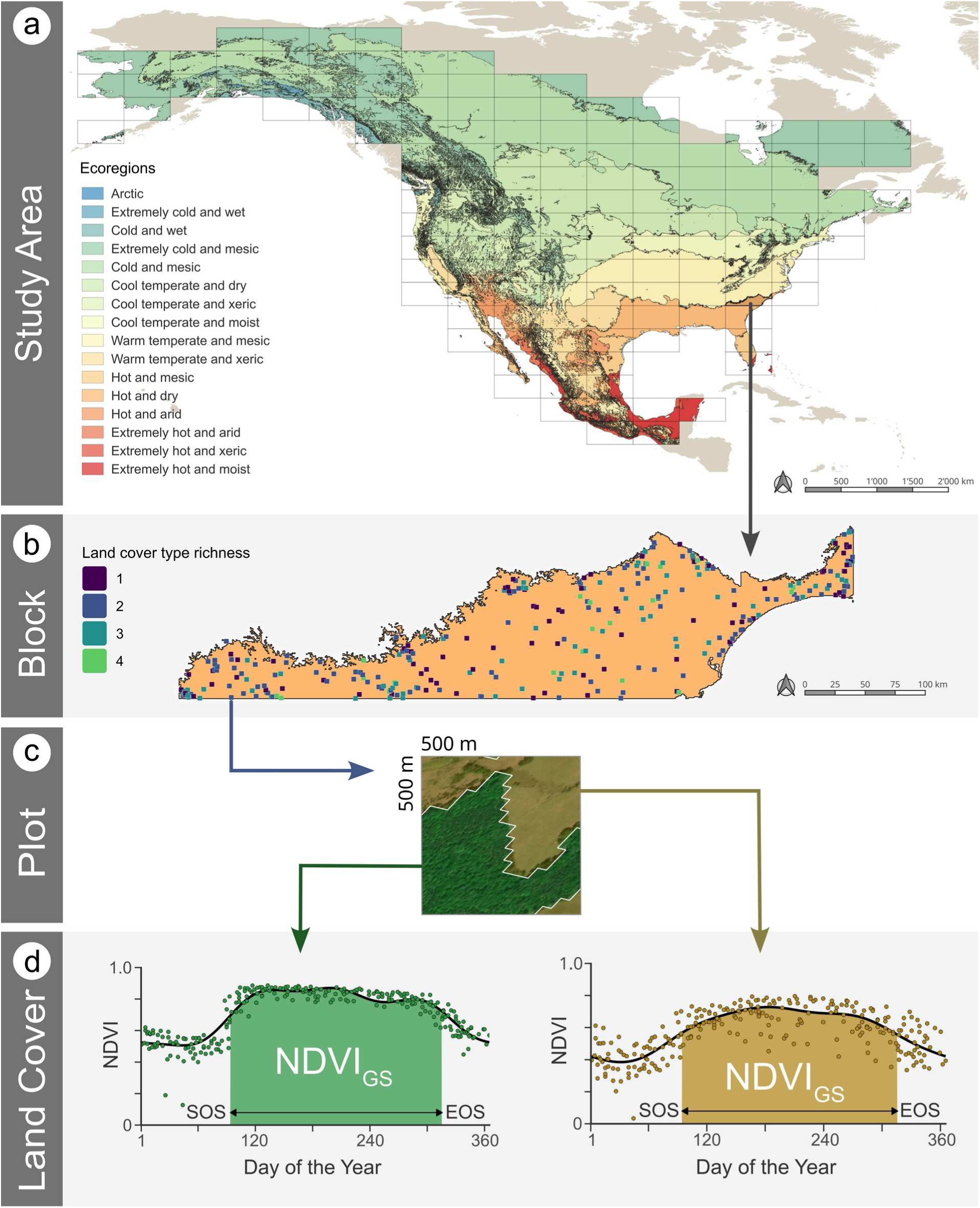
Study design. (**a**) Study area (continental North America) with ecoregions and 3° latitude x 6° longitude tiles, which together serve as blocks. (**b**) Example of a block with plots differing in land cover type richness (LCR). (**c**) Example of a plot with a LCR of 2, with (**d**) growing-season-integrated productivity indices (NDVI_GS_) determined for each of the two land covers, using harmonic analysis of time series (HANTS; see Methods). NDVI: normalized difference vegetation index; SOS and EOS: start and end of season.

First, we established that the landscape diversity-productivity relationships identified previously could be replicated using this new network of landscapes and using spatially more highly resolved Landsat satellite images (30 m resolution) instead of the previously used MODIS images (250 m resolution). Next, we asked whether net landscape diversity effects were disproportionally driven by a few LC types; to answer this question, we decomposed net landscape diversity effects into statistical selection and complementarity effects using the additive partitioning methods^20^. We then focused on inter-annual variation in landscape productivity. Specifically, we asked whether, under variable environmental conditions, the productivity of mixed-LC-type landscapes was temporally more stable than the one of single-LC landscapes, and whether such an increased temporal stability was mediated by contributions from different LC types in different regions, that is by a type of spatial insurance effect^21–23^.

## Results

### Landscape productivity is positively associated with land-cover-type richness

We quantified vegetation productivity by integrating Landsat-satellite-derived normalized difference vegetation indices (NDVI)^24^ over the growing season. The 15-year averages of these growing-season integrals (NDVI_GS_) were statistically significantly higher in mixed-LC landscapes than in single-LC landscapes (Fig. 2; positive net diversity effect (NE) on NDVI_GS_; t_42_ = 7.45, P < 0.001). Within LC-type mixtures, net diversity effects tended to increase further as more LC types were added (F_1,42_ = 3.3, P = 0.076). These findings are in line with previous, MODIS satellite data-based, analysis carried out for Switzerland^16^, and with a second analysis for North America^15^. This demonstrates that using data from either satellite yields comparable results, despite large differences in temporal (∼16x) and spatial (∼70x) resolution.

**Fig 2.**
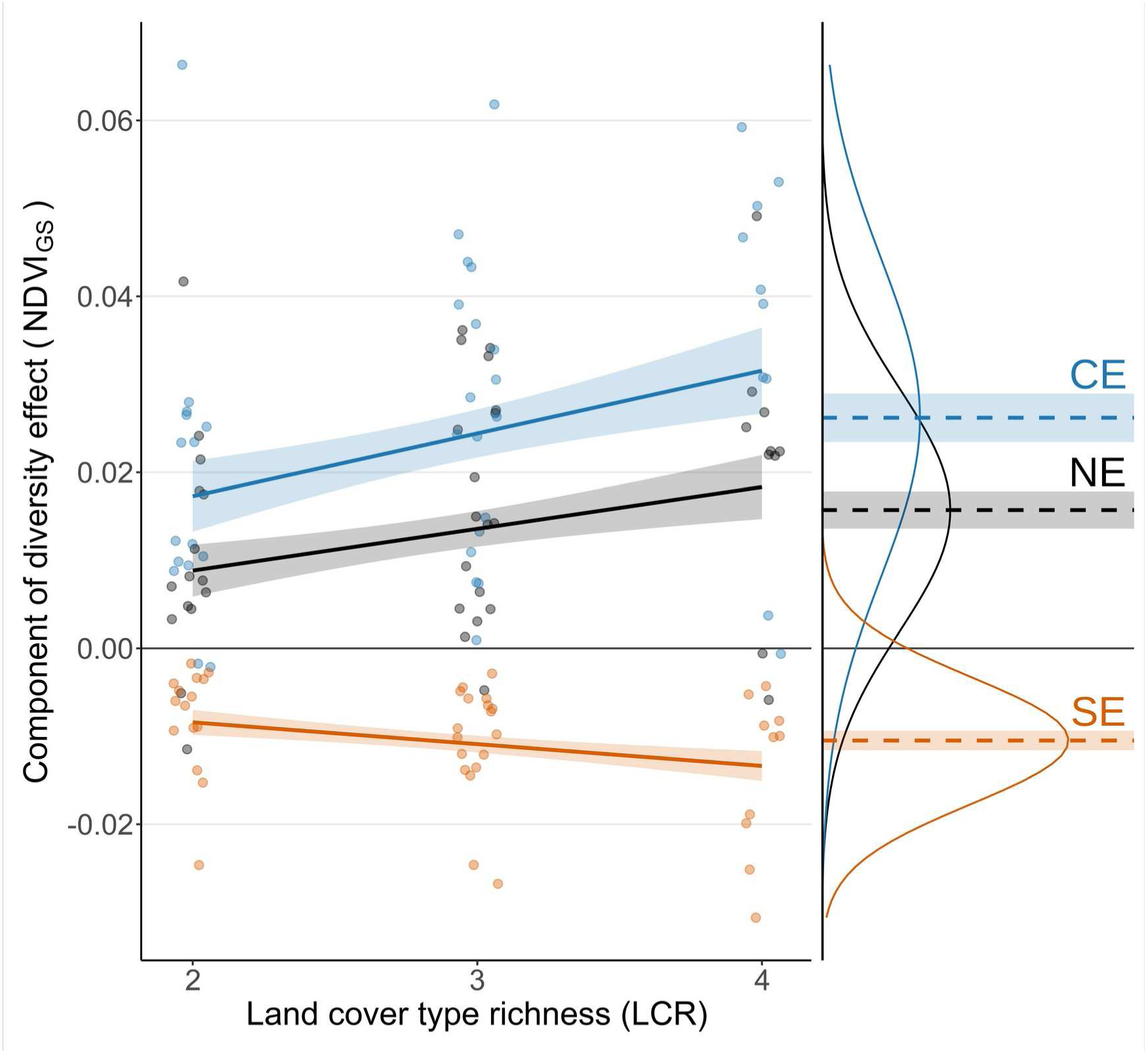
Additive partitioning of landscape diversity effect. Net diversity effect (NE, black), complementarity effect (CE, blue) and selection effect (SE, orange) in dependence of land-cover type richness (LCR). Lines and filled areas show model-predicted values (means±standard errors). Symbols show values for individual LC type compositions (n=43). The right panel show means and standard errors across all LC mixtures, with lines reflecting the standard deviation of the distribution formed by the 43 different compositions.

### Diversity effects are stable throughout growing season

A higher productivity in mixed-LC landscapes may result from a constant increase in the productivity of LCs in mixed landscapes, or from a lengthening of the period of active growth. The latter could occur, for example, when urban areas create heat that is redistributed into the surroundings, allowing for an earlier start of the season in these areas if vegetation growth is temperature-limited^25^. To investigate these contrasting possible causes of diversity effects, we divided the potential growing season into its interquartile range, which reflects conditions around peak season, and the lower and upper quartiles, which reflect conditions during the two shoulder seasons (Fig. 3), and then analyzed LCR-productivity relationships on NDVI_GS_ separately for each interval. The results indicate that the diversity effects observed in the three parts of the growing season are statistically indistinguishable (Table 1), i.e., there is no indication that the increased productivity in mixed-LC landscapes originates only from an extended growing season.

**Fig 3.**
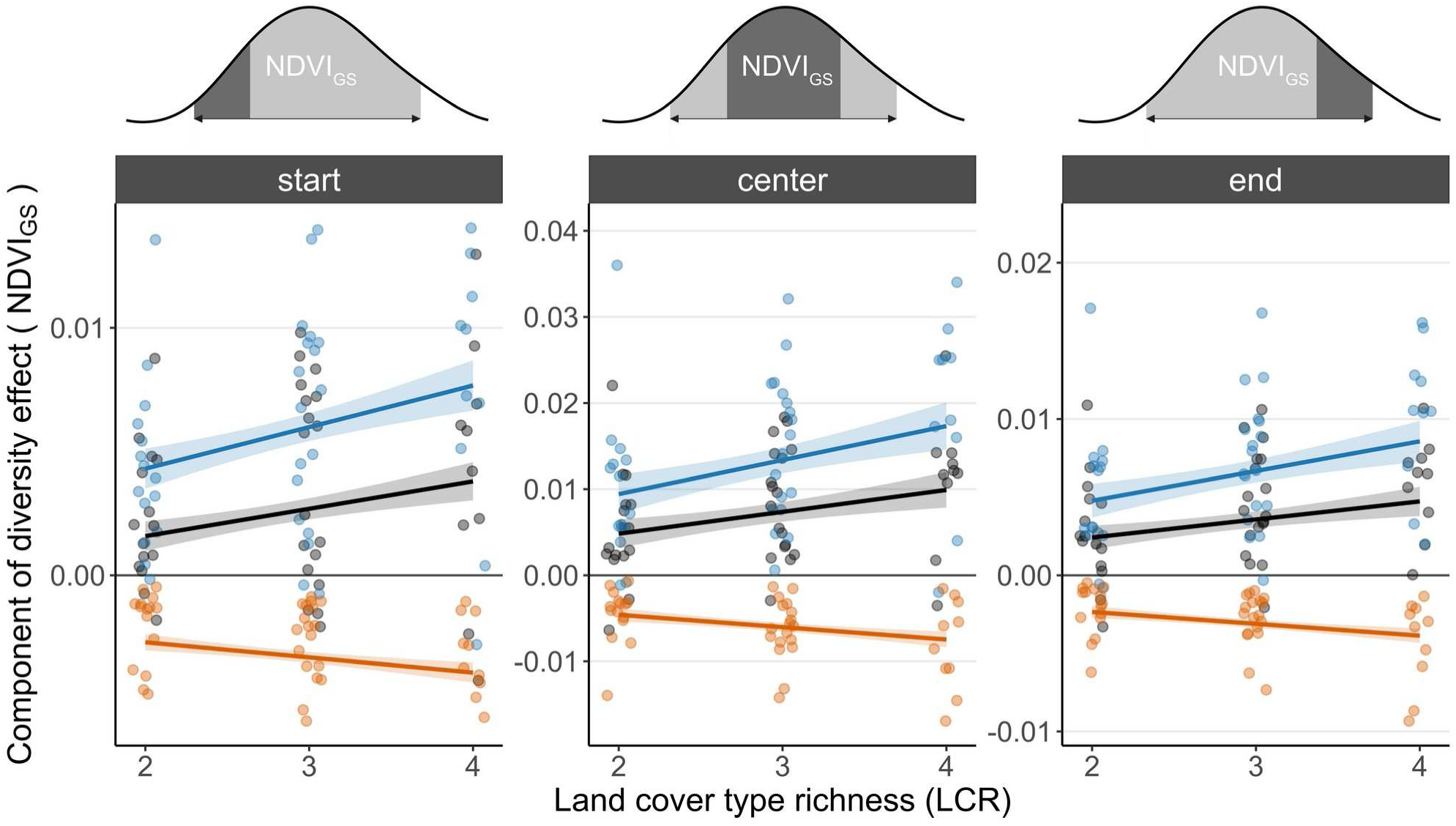
Seasonal dynamics of landscape diversity effect. Net diversity effect (NE, black), complementarity effect (CE, blue) and selection effect (SE, orange) in dependence of land-cover type richness (LCR). The growing season was divided into the first quartile (start), the second and third quartiles (center, i.e. the interquartile-range), and the fourth quartile (end). Center thus corresponds to the main growing season, and start and end to the shoulder seasons. Lines and filled areas show model-predicted values with standard errors, respectively.

**Table 1.** Statistical tests for net diversity effect (NE) and its complementarity and selection components (CE and SE) on productivity, quantified as growing-season integrated NDVI, during the early, peak, and late growing season. Top: tests for effects across all mixtures being positive (> 0). Bottom: tests for linear dependency of effects on land-cover type richness (LCR) within mixtures.

| Effect type | Component | Early season<br>(1st quartile) |  | Peak season<br>(inter-quartile range) |  | Late season<br>(4th quartile) |  |
| --- | --- | --- | --- | --- | --- | --- | --- |
| Component $> 0$ | NE | $t_{42} = 5.65$ | $P < .001$ | $t_{42} = 7.54$ | $P < .001$ | $t_{42} = 8.07$ | $P < .001$ |
| | CE | $t_{42} = 8.79$ | $P < .001$ | $t_{42} = 9.49$ | $P < .001$ | $t_{42} = 10.03$ | $P < .001$ |
| | SE | $t_{42} = -10.72$ | $P < .001$ | $t_{42} = -9.88$ | $P < .001$ | $t_{42} = -9.17$ | $P < .001$ |
| Linear<br>dependence of<br>component on<br>LCR | NE | $F_{1,41} = 1.62$ | $P = 0.211$ | $F_{1,41} = 4.58$ | $P = 0.038$ | $F_{1,41} = 4.86$ | $P = 0.033$ |
| | CE | $F_{1,41} = 3.25$ | $P = 0.079$ | $F_{1,41} = 6.24$ | $P = 0.017$ | $F_{1,41} = 7.96$ | $P = 0.007$ |
| | SE | $F_{1,41} = 3.47$ | $P = 0.067$ | $F_{1,41} = 4.98$ | $P = 0.031$ | $F_{1,41} = 6.35$ | $P = 0.016$ |

### Net diversity effects are driven by complementarity effects

The higher productivity of mixed LC landscapes is, by definition, driven by underlying productivity increases of one or several LC types. To characterize how productivity changes relative to the single LC situation (uniform land cover) are distributed among LC types, we applied Loreau and Hector’s additive partitioning method^20^. We found that the positive net diversity effects resulted from positive complementarity effects (CE), (t_42_ = 9.5, P < 0.001). In contrast, selection effects (SE) were slightly negative (t_42_ = -9.5, P < 0.001). Within LC mixtures, effect sizes of CE and SE increased with LCR (CE: F_1,42_ = 4.1, P = 0.05; SE: F_1,42_ = 4.0, P = 0.05). A positive CE indicates that, averaged across all LCs, there was a relative productivity increase relative to the respective single-LC landscapes (Fig. 2). A negative SE indicates that the relative productivity gains of the different LCs present in a mixed LC landscape decreases with the productivity of the respective single-LC landscapes. In other words, in mixed landscapes, LCs with generally lower productivity such as shrubland and wetland experienced stronger relative productivity gains than more productive LCs such as forest.

### Land-cover type diversity buffers large temporally variation in landscape functioning

Species-level biodiversity-ecosystem functioning experiments have shown that the productivity of mixed-species communities generally is temporally more stable than the one of less diverse communities^26,27^. To test whether conceptually comparable effects also occur at the landscape scale, we quantified the temporal variation of the productivity of each landscape as standard deviation of the inter-annual variation in NDVI_GS_ (σ_NDVIgs_; n = 14 years). The mechanisms that potentially reduce σ_NDVIgs_ in mixed LC landscapes are (1) contrasting seasonal dynamics of the component LC types (at species level, ref.^28^), or (2) interactions among LC types that reduce the temporal fluctuations in productivity within these through some other mechanism (e.g. species dominance, ref.^29^). We chose σ_NDVIgs_ as dependent variable because it quantifies variation at the plot-level in units of productivity, which is relevant from a landscape functioning perspective. Other commonly used metrics such as the coefficient of temporal variation (CV=σ_NDVI_/ *NDVI* ), or it’s inverse, express variation relative to each LC’s productivity, which is desirable for comparisons among LC types because it standardizes for productivity differences. However, under the null hypothesis of no diversity effects, the CV of a mixed-LC landscape will not be equal the mean CV of the respective single LC landscapes. The mathematical reason is that productivity values are additive, whereas relative changes in productivity are not.

We found that mean values of σ_NDVIgs_ did not depend on LCR (F_1,37_ = 0.24, P = 0.6) for LCR levels 1, 2 and 3 (LCR level 4 was not considered here because it did not occur in enough blocks). However, σ_NDVIgs_ was highly variable, and the range of this variability decreased with LCR (Fig. 4a). Specifically, the 95 % percentile of this distribution decreased with LCR, indicating that landscapes with extreme temporal fluctuations in productivity occur less frequently as LCR increases. Analyzing the temporal dynamics of productivity in all block ⨯ LC composition combinations, we found that the temporal variation in landscape productivity decreased most strongly relative to the temporal variation of the respective single LC landscapes when the productivity of the latter was most asynchronous (Fig. 4b). When we considered the asynchrony of the inter-annual productivity of the respective single-LC landscapes, expected values of σ_NDVIgs_ did no longer deviate from observations (Fig. 4c). Taken together, this indicates that the increased constancy of inter-annual variability was driven by the temporal asynchrony of the productivities of the different LC types.

**Fig 4.**
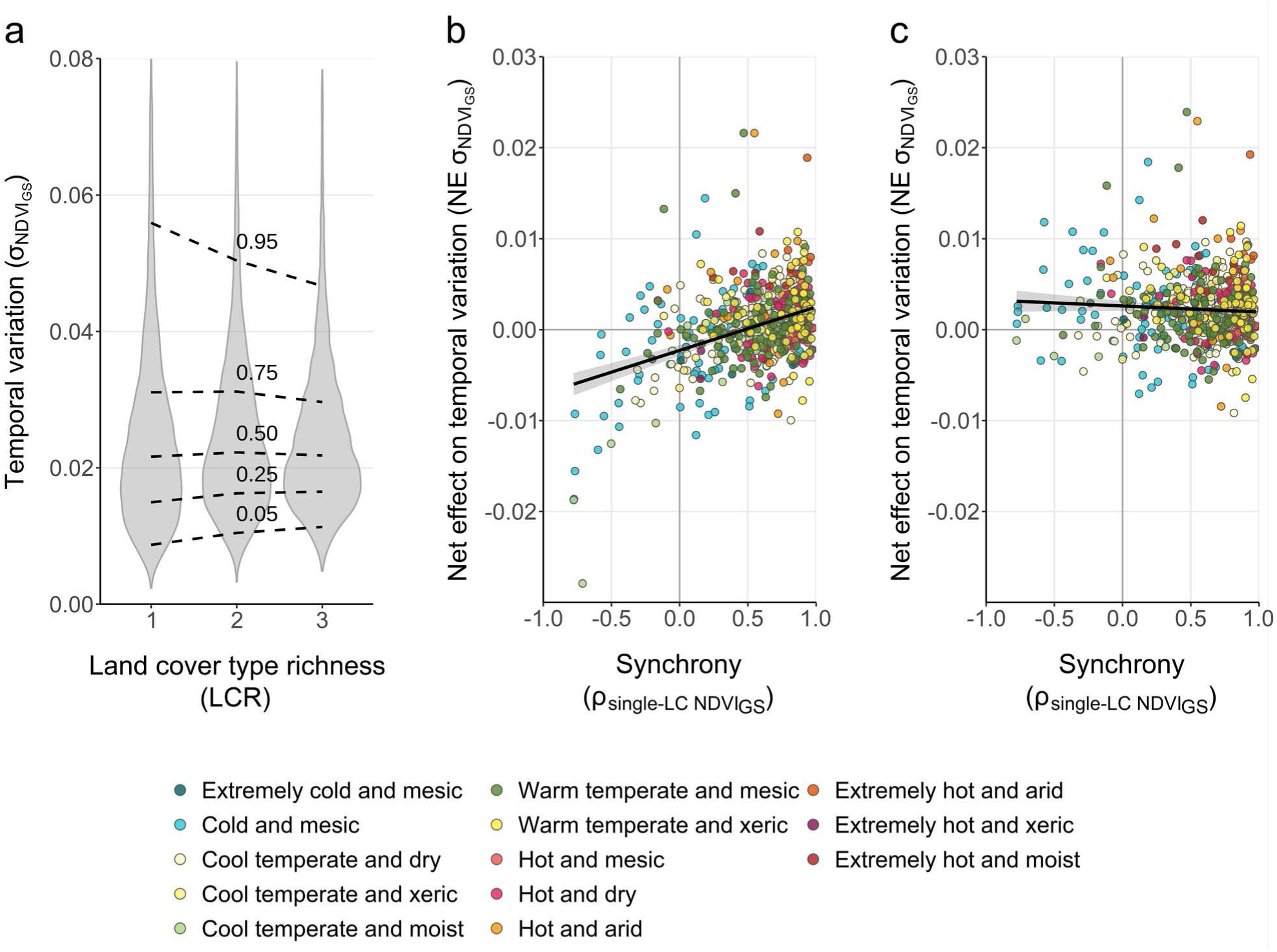
Temporal variability of productivity. (**a**) Inter-annual variation of productivity, quantified as standard deviation of NDVI (σ_NDVIgs_) in dependence of land-cover type richness (LCR). Lines show the respective quantiles of the distribution. (**b**) Net diversity effect on inter-annual variation of productivity (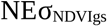) in dependence of inter-annual correlation of the respective single-LC landscape values (dots: binary LC mixtures in the different ecoregions). (**c**) Net diversity effect on inter-annual variation as in (b), but first correcting for the asynchrony of single LC landscape values in the calculation of the expected value. (b) and (c) together show that the net effects are driven by the asynchrony of interannual variation in NDVI among the different land covers present in a landscape.

### Drought is important temporal modulator of productivity

What are the factors that drive inter-annual variation in NDVI_GS_ at the level of the studied 25-ha landscapes? As one possibility, this variation is caused by local-scale stochasticity. In this case, the total productivity of larger areas of land would be stable because the productivity variation of individual landscape patches would be uncorrelated and therefore average out. Another, contrasting possibility is that the temporal variation is driven by specific external drivers that vary only at large spatial scales. In this case, the productivity of nearby landscape patches would fluctuate in unison, at least when they are of the same LC. This temporal variation would not average out at larger scales but would persist. Candidate drivers of such correlated variation in productivity include climatic factors (e.g. temperature, precipitation), but biotic causes such as larger-scale pest outbreaks may also be important. For our study, we postulated that drought was the most significant factor causing productivity variation.

Drought occurs when there is an imbalance between water inputs to an ecosystem, such as precipitation, and losses, such as evapotranspiration and drainage^30^. This balance also depends on water retention, which in turn depends on soil texture. Drought therefore is complex to model and only partly captured by single variables such as temperature and precipitation. We therefore instead used soil moisture time series from a global climate model re-analysis by the European Centre for Medium-Range Weather Forecasts, which includes land surface hydrology^31^. We reasoned that not all water in the soil profile is plant-available, and that a lack of water prior to the start of the growing season could have legacy effects on productivity. We explored different possibilities and found that soil moisture in the first meter of the soil profile, averaged over the growing season plus a few months prior to its start, was a good predictor of drought effects (see Methods); however, the exact choice of these parameters was not critical. Specifically, we used the time-integrated water deficit below a threshold as predictor of drought effects on landscape productivity (Fig. 5). This drought index explained up to about 80 % of the inter-annual variation in a block’s average NDVI_GS_, suggesting that drought is the single most important modulator of productivity, at least in warm and temperate dry regions. For example, a large fraction of productivity variation was explained by this index in the Great Plains, whereas much less variation could be explained in Arctic and cold regions such as northern Canada and Alaska as well as the central Rocky Mountains (Supplementary Fig. 1).

**Fig 5.**
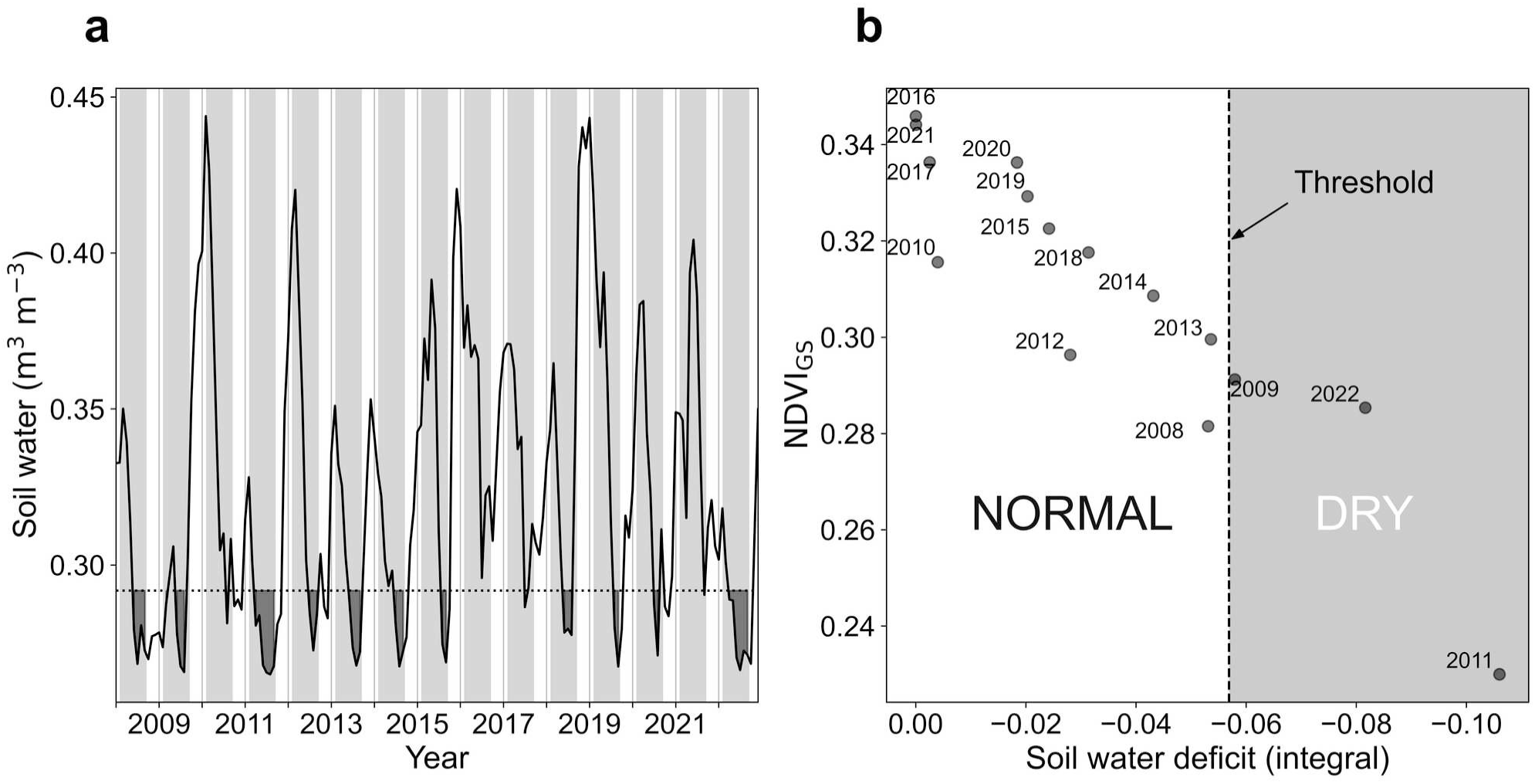
Soil water deficit determined for an exemplary block. (**a**) Time-course of monthly volumetric soil water (0-100 cm) (solid line). The dotted line is the soil moisture threshold under which growth is considered water-limited. Light gray areas show the potential growing season, and dark gray areas below the moisture threshold quantify the integrated soil water deficit, which we use as index of drought severity. (**b**) Productivity (NDVI_GS_) in dependence of drought severity index. The vertical dashed line divides years into normal and dry conditions (see Methods for details).

### Spatio-temporal selection effects are small

The absence of positive SE demonstrates that the diversity effects we found are not caused by individual, particularly productive, LC types at the continental scale (Fig. 2). However, the partitioning analysis applied so far was based on spatio-temporal averages across 15 years and 290 blocks. Selection effects may nevertheless be important at other scales or in particular years. If such effects are driven by different LC types in different areas or years, these local selection effects could manifest as complementarity effects when analyzing average values for each composition. This phenomenon is also known as "insurance effect" because a high landscape diversity ensures that under contrasting conditions at least one (but not necessarily always the same) of the many LC types ensures high landscape functioning levels. To test for this possibility, we applied a spatio-temporal partitioning method^23^ that extends the original additive partitioning method with terms accounting for selection effects arising across space and time.

First, we used the individual 25-ha landscapes as unit of replication, and partitioned all variation across the ∼50,000 landscapes of the network. We found a positive total CE (Fig. 6), which is in line with the results from the additive partitioning analysis (Fig. 2). The remaining term, the total selection effect, was negative (Fig. 6), again reflecting results from the additive partitioning. We then decomposed the total SE into components reflecting spatio-temporal variation, and residual terms that account for productivity differences of the LC types, so called average selection effects and non-random overyielding (see Table 2 and ref.^23^ for details). We found that the spatial selection effect was positive (Fig. 6, see Supplementary Table 1), i.e., the LC types that performed particularly well differed between locations. Interestingly, despite drought being such a strong modulator of productivity, the classification of years into dry and normal years did not explain temporal differences in higher productivity with respect to selection effects, i.e. the temporal and spatio-temporal selection effects were negligible.

**Fig 6.**
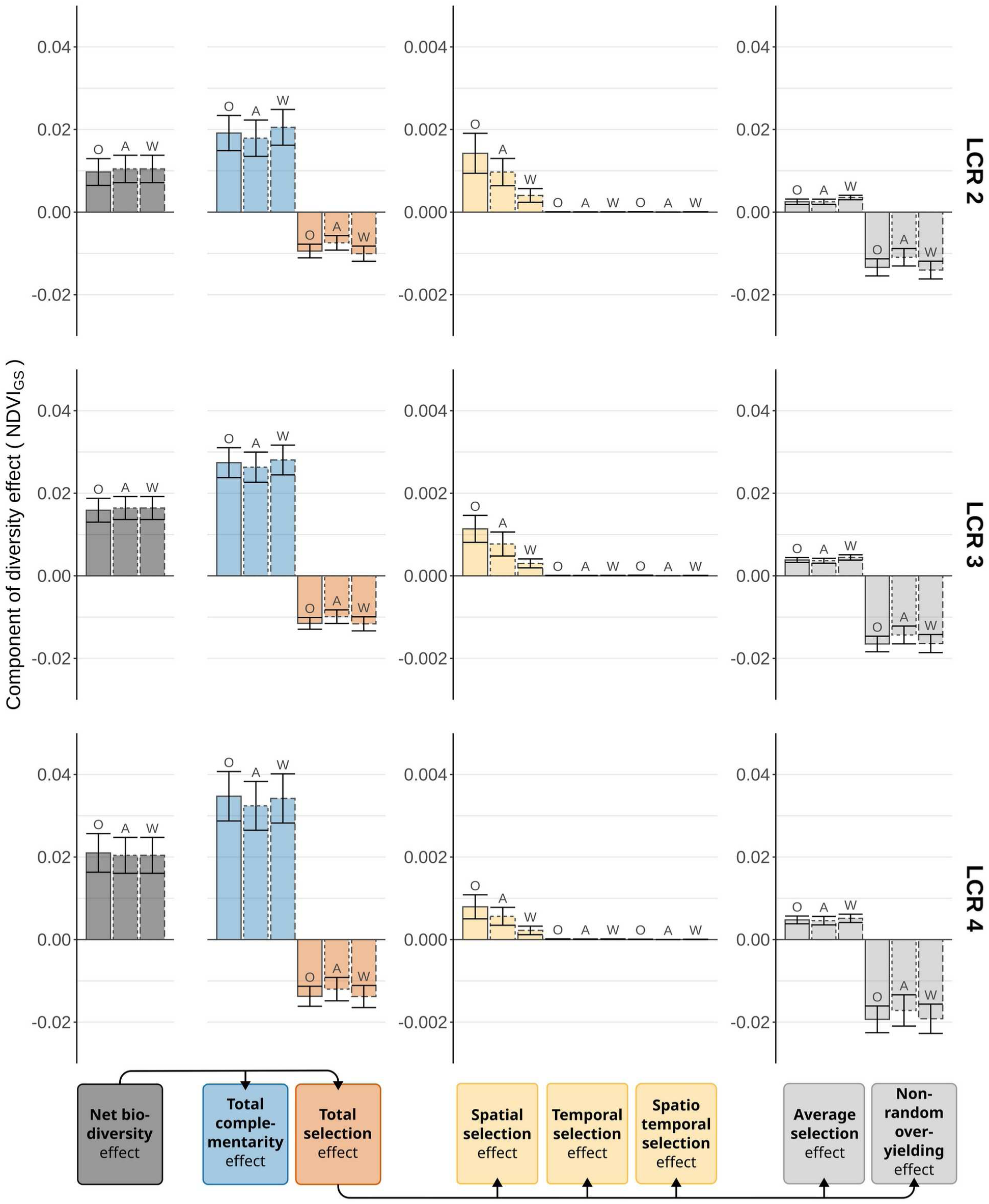
Spatio-temporal partitioning of net diversity effect. The spatio-temporal net diversity effect values for NDVI_GS_ were partitioned into a total complementarity and a total selection effect (left; spatio-temporal averages). The total selection effect was then further partitioned into a temporal, a spatial, and a spatio-temporal component (center), and residual terms (right; average selection effect and non-random overyielding). This partitioning was performed for the total variation observed across all ecoregions and blocks (O); for the variation between ecoregions (A); and for the variation within ecoregions (W) (see Methods for details). Results are shown for all LC-type richness (LCR) levels 2, 3 and 4.

**Table 2.**
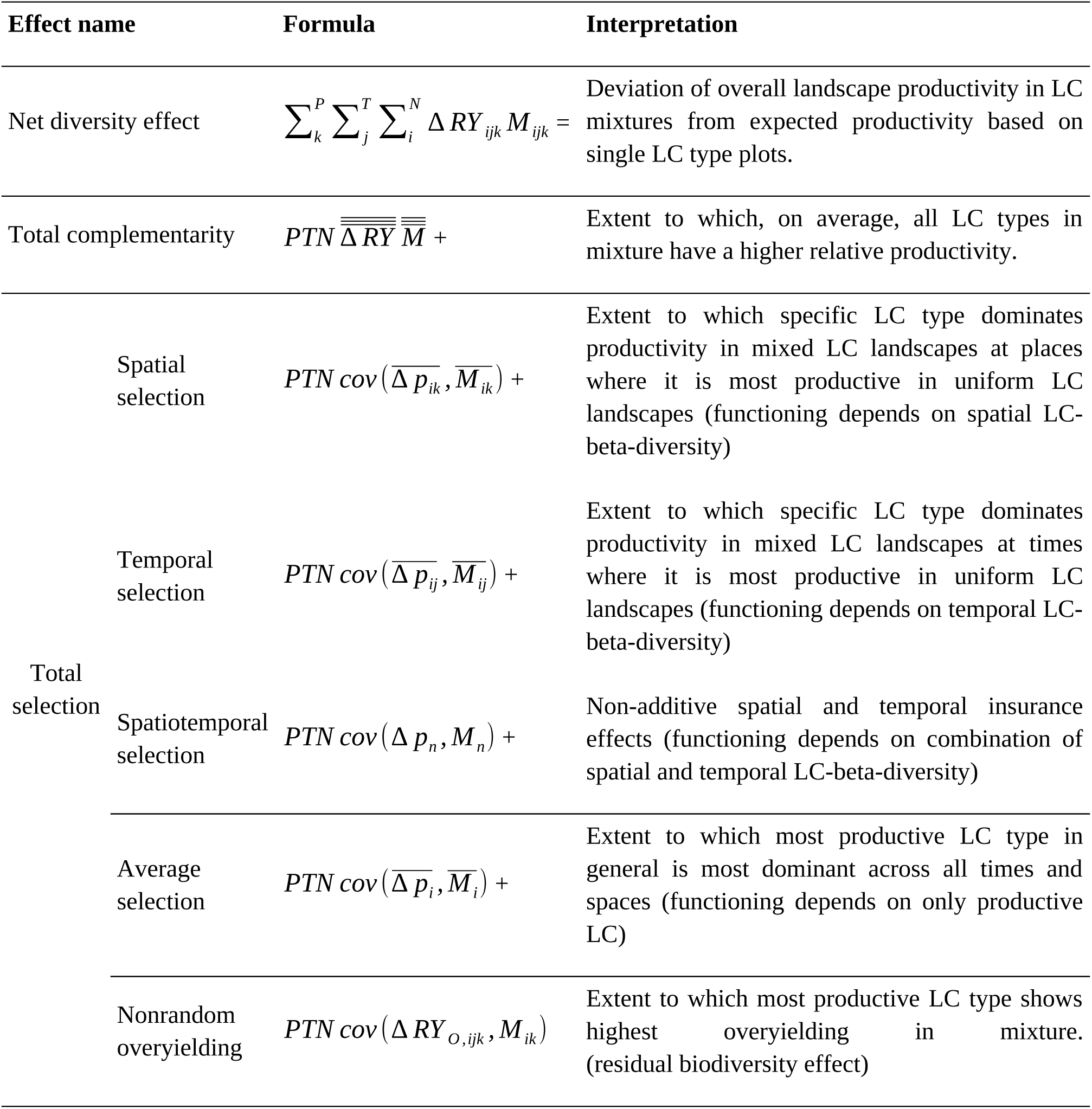
Effect parts, formula and suggested interpretation for spatio-temporal partitioning of effects (for details, see ref.^23^). Place *k* in total places *P*, time *j* in total times *T* and land cover (LC) type *i* in all LC types *N* in specific composition. Δ*RY = RY_O_ - RY_E_* with *RY_O_ = Y_O_/M* where *M* is the observed yield in uniform single-LC landscapes and Y_O_ the observed yield in mixed-LC landscapes. Overall, subscript *O* refers to observed and *E* to expected values. Δ*RY_O,ijk_ = RY_O,ijk_ – p_O,ijk_* with *p_O,ijk_* as the observed relative productivity of LC type *i* at time *j* and place *k*. Therefore, Δ*p_O,ijk_ = RYE_,ijk_ – p_O,ijk_*.

| Effect name | Formula | Interpretation |
| --- | --- | --- |
| Net diversity effect | $\sum_k^P \sum_j^T \sum_i^N \Delta RY_{ijk} M_{ijk} =$ | Deviation of overall landscape productivity in LC mixtures from expected productivity based on single LC type plots. |
| Total complementarity | $PTN \overline{\overline{\overline{\Delta RY}}} \overline{\overline{\overline{M}}} +$ | Extent to which, on average, all LC types in mixture have a higher relative productivity. |
| Total selection | Spatial selection<br>$PTN cov(\overline{\Delta p_{ik}}, \overline{M_{ik}}) +$ | Extent to which specific LC type dominates productivity in mixed LC landscapes at places where it is most productive in uniform LC landscapes (functioning depends on spatial LC-beta-diversity) |
| | Temporal selection<br>$PTN cov(\overline{\Delta p_{ij}}, \overline{M_{ij}}) +$ | Extent to which specific LC type dominates productivity in mixed LC landscapes at times where it is most productive in uniform LC landscapes (functioning depends on temporal LC-beta-diversity) |
| | Spatiotemporal selection<br>$PTN cov(\Delta p_n, M_n) +$ | Non-additive spatial and temporal insurance effects (functioning depends on combination of spatial and temporal LC-beta-diversity) |
| | Average selection<br>$PTN cov(\overline{\Delta p_i}, \overline{M_i}) +$ | Extent to which most productive LC type in general is most dominant across all times and spaces (functioning depends on only productive LC) |
| | Nonrandom overyielding<br>$PTN cov(\Delta RY_{O,ijk}, M_{ik})$ | Extent to which most productive LC type shows highest overyielding in mixture. (residual biodiversity effect) |

Next, we asked whether the spatial selection effect was predominantly driven by spatial variation among biogeographic regions, i.e. by variation at a large scale, or whether it was driven by variation within these regions. To investigate these possibilities, we repeated the spatio-temporal partitioning using either average values for each combination of LC type and biogeographic regions, or by performing the partitioning within each biogeographic region and then averaging the results. Our results indicated that the overall spatial selection effect was driven by both of these components.

## Discussion

Applying the additive partitioning method to net landscape diversity effects, we found evidence that landscape functioning benefits from the presence of a high diversity of different LC types. Complementarity effects (CE) were overall positive, indicating that interactions among LC types were on average beneficial for the LCs participating in these interactions. Also, selection effects (SE) were on average negative, indicating that more productive LCs did not benefit disproportionally. In other words, landscape diversity effects were not driven by single highly productive LC types. These statistical patterns of relative yields are remarkably similar to what has been found in plot-level BEF experiments, especially in the ones that ran for several years^32–34^. In plot-scale BEF experiments, productive, dominant, species typically gain in cover as the community establishes after sowing, which contributes to a high community biomass^35,36^. Such shifts could in principle also occur among LC types, for example when a forest expands into adjacent grassland. Such encroachment is observed in many regions globally, and may contribute to productivity increases^37^. However, in our study the areal fractions which the different LC types occupied in a landscape remained fixed. The phenomenological similarity in net diversity effects and the underlying CE and SE therefore appear all the more remarkable.

The specific biological mechanisms that underlie these effects at the landscape level are not well studied to date^7^. However, there is evidence that landscape diversity can be functionally important, even though the underlying mechanisms are not yet well understood. As one possibility, a high landscape diversity begets a high local species diversity, which in turn promotes landscape functioning through the species interaction mechanisms established in plot-scale BEF experiments^38^. This indirect effect of landscape diversity could occur when additional niche space is created at boundaries between different ecosystem types and in turn leads to higher species richness, or when local species richness is increased due to meta-population and meta-community processes^39–41^. A contrasting possibility is that different ecosystems interact through mechanisms that are largely independent of local species richness, and that these interactions are on average non-neutral and thereby systematically promote landscape functions such as productivity. For example, nutrients are commonly redistributed across ecosystem boundaries, which often supports primary production in the more nutrient-limited system^21,42^. As another example, landscapes comprising a mixture of forest and grassland often are cooler than the average of the corresponding single land-cover landscapes^43^. This temperature change likely occurs because differences in the surface properties between ecosystem types cause atmospheric turbulence, which promotes heat transport away from the land surface. This effect could be particularly beneficial at high temperatures. Overall, this indicates that landscape diversity has the potential to affect local climate, which is relevant for productivity.

Focusing on the spatio-temporal variation of these patterns, we found that spatial selection effects were important. In other words, productive LCs supported local landscape productivity, but the relevant LCs differed by location. Overall, a high diversity of LC types thus presents a type of “spatial insurance” with respect to landscape functioning. However, it is important to note that a component of the local selection effects will manifest in a positive total complementarity effect when these selection effects are on average positive.

Interestingly, the temporal component of this variation was quantitatively unimportant. Our data showed asynchronous inter-annual dynamics of the different LC types, and this asynchrony was in part related to drought. Asynchronous responses of the functions of different ecosystem types to stress are common. For example, dry tropical forest and rain forest show asynchronous phenology under El-Niño and seasonally-induced drought in Hawaii^44^. Another example are temperate and tropical forest edges, which were found to be more susceptible to drought but also recover faster than interior forest areas^45,46^. In our study, such asynchronous temporal dynamics did not manifest as temporal selection effects.

We showed evidence that LC-type diversity is positively associated with landscape-level productivity, and that the types of ecosystems that support system-level productivity vary spatio-temporally. To date, the primary way to leverage diversity in the management of ecosystem services is to promote local species richness, for example through conservation measures in natural and semi-natural habitats, or by increasing diversity in agricultural production systems. Our study suggests that an additional possibility is to promote diversity at a higher level of ecological organization, specifically by maintaining a complex land-cover structure. From an applied perspective, this is interesting because on a large part of the terrestrial surface, land cover is determined by human land-use and management^47^. However, to effectively do so, the specific biological mechanisms and the spatial scale at which land-cover diversity exerts the largest positive effects need to be better understood.

## Methods

### Study design

To relate landscape functioning to landscape diversity, we established a large number of ∼25 ha study plots spread across North America. Specifically, we adopted a quasi-experimental design based on experimental design principles widely applied in plot-scale biodiversity-ecosystem functioning studies^18^. To separate the diversity effects of interest from effects of environmental variation at large spatial scales, we stratified North America spatially and according to environmental variables, similarly to how the field sites of experimental studies are commonly divided into blocks and the treatments of interest then replicated within these blocks. We divided North America into 16 ecoregions defined in the Global Environmental Stratification dataset^19^. We did not consider some arctic areas due to their low plant productivity and because these high-latitude areas are poorly covered by the satellite remote-sensing products we used as proxies of landscape functioning. To control heterogeneity within ecoregions, we further divided these into 3° latitude ⨯ 6° longitude blocks (Fig. 1).

### Landscape diversity

As measure of landscape diversity, we used the number of different land-cover (LC) types found in each landscape plot (LC type richness, LCR). The LC types were based on the Commission for Environmental Cooperation’s North American land monitoring system map^17^ for 2015 (30 m spatial resolution). We aggregated the original 19 LC classes into 6 broad, ecologically distinct classes: agriculture, forest, grassland, shrubland, wetland, and urban areas. In addition, topographic variables (slope, gradient and elevation) were calculated from the digital elevation model TanDEM-X^48^ which has a spatial resolution of 90 m. We used this topographic data to create LC diversity gradients that were orthogonal to topographic gradients that likely also affected landscape productivity. Specifically, we selected such a subset of plots using simulated annealing, a probabilistic subsampling technique (see ref.^15^ for details). The selection follows an orthogonal design, ensuring that an equal number of replicates is selected for each LC composition level realized in a block. Specifically, we selected 20 replicates of each realizable composition within a block.

### Landscape functioning

As a measure of landscape function, we used the Normalized Difference Vegetation Index (NDVI), which is commonly used as proxy of vegetation productivity ^49,50^. NDVI was calculated based on Landsat satellite data (years 2008 to 2022), using Google Earth Engine^51^. NDVI data from Landsat has a temporal resolution of 16 days, which is low compared to the rate of vegetation development, in particular during the shoulder seasons. To better cover this seasonal dynamic, we therefore combined data from Landsat 8, 7 and 5. We excluded acquisitions potentially affected by clouds by excluding areas under or near clouds (500 m buffer around areas with a low likelihood of cloud shadow), and removed blocks for which only limited data was available. This resulted in a data set encompassing 290 blocks with a total of 50,682 landscapes representing 49 different LC type combinations. We then generated 15-year NDVI time series for each LC within each landscape separately, and modeled these data as sum of three harmonics using the HANTS method, which is insensitive towards the type of outliers that occur frequently with these satellite data ^16,52^. The resulting model of vegetation phenology has annual periodicity, and was used to determine the growing-season integral of NDVI (NDVI_GS_) by integration from start to end of the growing season (SOS and EOS). SOS and EOS had previously been determined by applying the NDVI-ratio method to a random sample of 1,000,000 vegetation index values (250 m ⨯ 250 m pixels from MODIS satellite instrument) per ecoregion. We then used the 25 % and 75 % percentiles of the SOS and EOS distribution as SOS and EOS of the respective ecoregion ^15^. The rationale underlying this choice is that this interval approximates the period during which meaningful vegetation activity is potentially possible, yet SOS and EOS estimation remain robust in light of the inherently high variability of NDVI-ratio results. All NDVI values were further scaled by dividing these by the average value of a large random sample of NDVI values per ecoregion, the rationale being to normalize all calculations to the average productivity of the region.

### Additive partitioning

The net diversity effect (NE) was calculated as NDVI_GS_ of each landscape, minus the mean NDVI_GS_ of the respective single-LC landscapes (average values in respective block). NE was then decomposed in complementarity (CE) and selection effects (SE) following Loreau & Hector^20^. CE and SE are based on relative yields (*RY*). Because NDVI is a measure of density, i.e., an index of productivity per unit land area, we first scaled NDVI_GS_ by the respective land area fraction *x*. *RY* of each LC in each mixture was thus calculated as 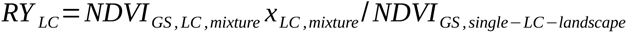. CE and SE were determined at the block level, and then values first aggregated to the ecoregion and then averaged across ecoregions.

### Inter-annual variation in productivity

The temporal variability of the productivity of each landscape was quantified as standard deviation σ of the inter-annual variability of NDVI_GS_ (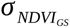). These calculations were done excluding landscapes with a LCR of four because in many blocks no corresponding mixtures existed. We then used the NDVI_GS_ times series of the uniform single-LC landscapes, averaged at the block level, to determined the correlation of the time series between all LC pairs. Specifically, we determined the synchrony of their time course. Next, we determined the net diversity effect (NE) on the inter-annual variation, for all binary LC mixtures, 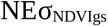, in two different ways: first, we used the mean 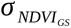 of the uniform single-LC landscapes as expected value, i.e., for a LC mixture of A and B, 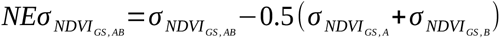; second, we considered the covariance in the calculation of the expected value, so that 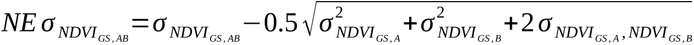. The latter value is larger than the former when the NDVI values of the two LC don’t vary fully synchronously.

### Drought index

We retrieved volumetric soil water in the soil top 100 cm for each landscape from ERA5-Land monthly averaged data^31^. Then, separately for each block, we calculated a drought index (DI) by defining a threshold under which we expected that plant productivity would be reduced, and integrated this water deficit over a period starting a few months prior to the start of the season (SOS) to ¾ of the growing season (EOS). The number of months prior to SOS and the soil moisture threshold where varied to find the values that best predicted the observed inter-annual variation in the block’s median NDVI_GS_.

### Spatio-temporal partitioning

Isbell et al.^23^ extended the original additive partitioning method in order to partition spatial and temporal components of net biodiversity effects (NE). Table 2 summarizes the terms into which NE is decomposed (see original paper for details). Here, we focus on the spatial selection effect, the temporal selection effect, and the spatio-temporal selection effect. Note that in the original publication these terms are referred to as “insurance effects”. However, since what is partitioned are selection effects, we follow the suggestion of the original authors and refer to them as spatial and temporal selection effects^53^.

To test whether the spatial selection effects were predominantly driven by small-scale or by large-scale variation, we repeated the spatio-temporal partitioning considering entire ecoregions as places in space, and eliminating variation between ecoregions and considering blocks within ecoregions as spaces. The former was achieved by analyzing data aggregated at the level of ecoregions, and the latter by partitioning variation among blocks for each ecoregion separately, and then averaging results across ecoregions.

Given that drought was a strong modulator of NDVI_GS_, and therefore likely the functionally decisive component of time, we classified years into dry and normal years, separately for each block, and used this classification instead of calendar year as measure of time. Analyses of long-term climate records have shown that approximately each 6^th^ to 7^th^ year can be considered as exceptionally dry^54^.We therefore sorted the years by drought index, and classified the years with the 16 % highest values as dry. The spatio-temporal partitioning requires that all contributing LC type occur at all places and under dry and normal conditions. To meet these requirements, we performed the partitioning on a subset of the data (243 blocks comprising of a total of 42,742 landscapes).

### Statistical analysis

All data were analyzed using linear mixed-effect models summarized in analysis of variance (ANOVA) tables, using ASReml-R package^55^ (VSN International, Hemel Hemsted, UK). Blocks and LCR were fixed effects, and LC composition was a random effect, i.e., the different LC compositions were considered the statistical replicates in the LCR gradient.

## Acknowledgments

This project was funding by a grant of the University of Zurich Research Priority Program Global Change and Biodiversity (URPP GCB) to PAN and RF. SL received additional funding from the European Union’s Horizon 2020 research and innovation programme under the Marie Sklodowska-Curie grant agreement No. 847585.

## Author contributions

PAN, FA and RF conceived the idea. PAN, ES and RF acquired funding. SL implemented the study and analysed the data, primarily supported by PAN. SL was supported with data interpretation by ES and FI during research stays. SL and PAN wrote the first draft. ES, FA, FI, and RF contributed to the final manuscript.

### Boxes

#### Box 1. Glossary

**Landscape diversity** | landscapes encompass a variety of landforms, environmental conditions, and habitat types. Therefore, diversity in a landscape can be quantified in different ways, including species diversity (→**gamma diversity** across different habitats), and as number of distinct →**land-cover** types (→**alpha diversity** of **land cover** types), which is central to the present work.

**Alpha diversity** | the diversity of components within a relatively uniform system. Examples include the number of species within a specific type of plant community and the number of land-cover types within a single landscape patch.

**Gamma diversity** | the diversity of components across a heterogeneous set of systems – such as the total number of species found in various habitats within a landscape.

**Land cover |** describes the physical and biological characteristics of a given area’s surface. Types of land covers include different vegetation types, such as forests, grassland, wetlands, and agricultural areas, as well as non-vegetated and anthropogenic areas, such as open water and urban areas.

**Normalized difference vegetation index (NDVI )** | a widely used remote sensing metric for estimating standing green biomass. It is calculated using reflected red and near-infrared light and is strongly correlated with primary productivity.

**Additive partitioning** | statistical method developed by Loreau & Hector^20^ to decompose net biodiversity effects (NE) into **complementarity effects** (CE) and →**selection effects** (SE). This method is based on deviations of →**relative yields** (RY) from the null expectation of RY = 1/n in an even mixture of n components. Under the null hypothesis of no diversity effect, ΔRY equals zero.

**Relative yield (RY)** | measure of the productivity of a component in a mixture relative to its productivity when it exists alone (e.g. a species grown as a monoculture, or a land-cover in a landscape dominated by a single land cover type).

**Complementary effect (CE)** | component of the net biodiversity effect (NE), derived using the →**additive partitioning** method. CE quantifies the average benefit that components gain from being in a mixture, relative to their performance when alone. CE is proportional to the average ΔRY across all components.

**Selection effect (SE)** | component of the net biodiversity effect (NE), derived using the →**additive partitioning** method. SE quantifies the extent to which each component’s relative benefit from being part of a mixture is related to its productivity when it is the only part of the system. SE is proportional to the covariance between ΔRY and productivity when alone.

**Spatio-temporal partitioning** | extension of the →**additive partitioning** method that incorporates spatial and temporal variation in the contributions of different components to the net diversity effect (NE). Developed by Isbell et al. ^23^, it is conceptually grounded in the →**insurance hypothesis**.

**Insurance hypothesis** | proposes that systems containing a higher number of different components are more resilient, as they are more likely to include components that maintain system functioning under changing environmental conditions. This concept applies to both spatial and temporal environmental variability.

## Supplementary Material

**Supplementary Fig 1.**
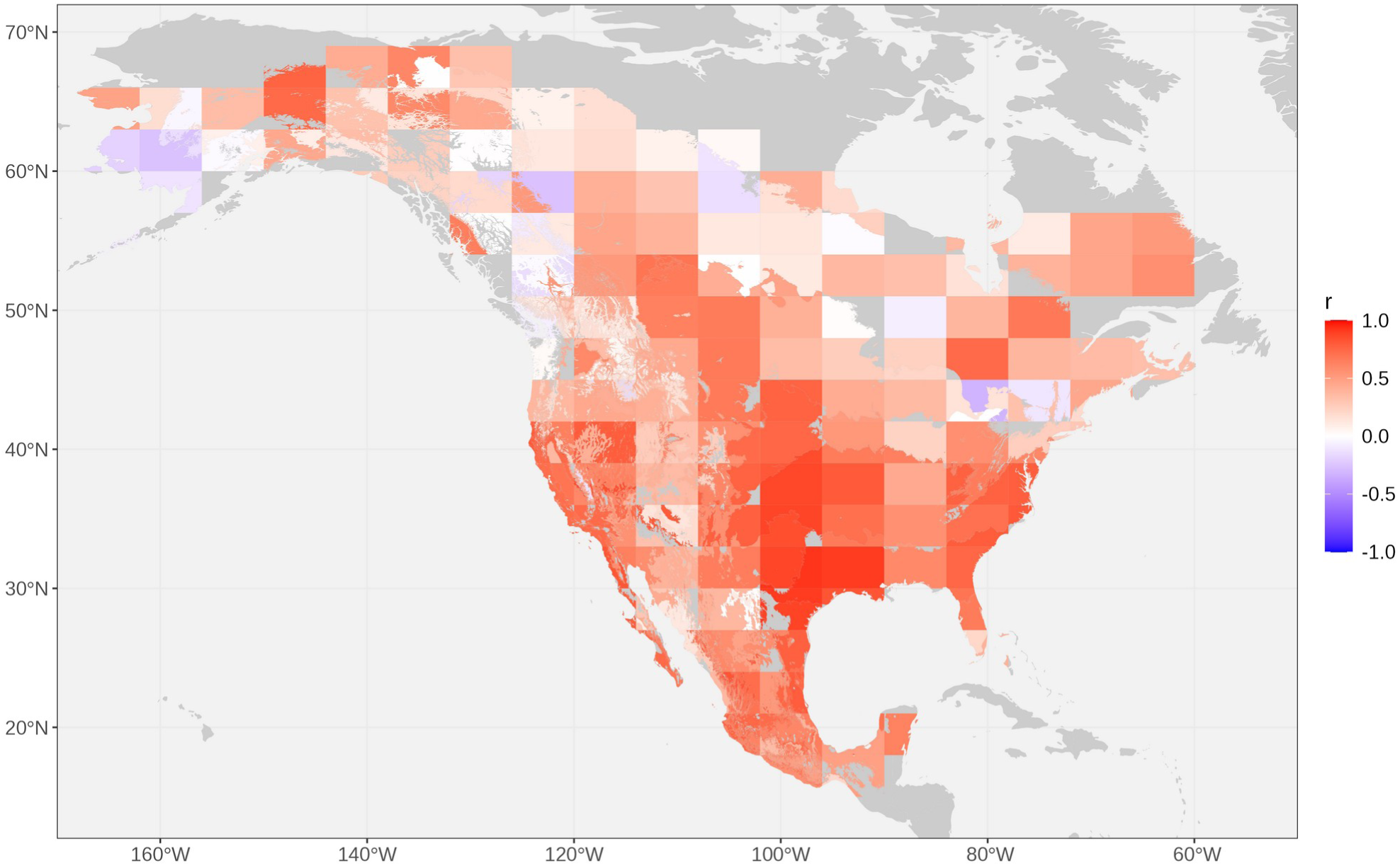
Correlation coefficient for drought index (integral under threshold, see Methods) and NDVI_GS_ mean for all plots in block.

**Supplementary Table 1.** Spatiotemporal partitioning results expressed as percentage of net diversity effect (NE). O = overall (blocks represent different places), A = across ecoregions (ecoregions represent different places), W = within ecoregion (blocks represent different places within each ecoregion).

| LCR | Group | Net diversity (%) | Total complementarity (%) | Total selection (%) | Spatial selection (%) | Temporal selection (%) | Spatio-temporal selection (%) | Average selection (%) | Non-random overyielding (%) |
| --- | --- | --- | --- | --- | --- | --- | --- | --- | --- |
| 2 | A | 100.00 | 171.14 | -71.14 | 9.27 | 0.04 | 0.03 | 24.07 | -104.56 |
| 2 | O | 100.00 | 196.98 | -96.98 | 14.65 | 0.08 | 0.11 | 25.92 | -137.74 |
| 2 | W | 100.00 | 196.22 | -96.22 | 3.87 | 0.07 | 0.07 | 33.93 | -134.16 |
| 3 | A | 100.00 | 160.24 | -60.24 | 4.69 | 0.04 | 0.02 | 22.24 | -87.23 |
| 3 | O | 100.00 | 172.54 | -72.54 | 7.16 | 0.05 | 0.08 | 24.12 | -103.95 |
| 3 | W | 100.00 | 170.86 | -70.86 | 1.83 | 0.06 | 0.04 | 27.11 | -99.90 |
| 4 | A | 100.00 | 158.77 | -58.77 | 2.76 | 0.05 | 0.01 | 22.50 | -84.10 |
| 4 | O | 100.00 | 165.39 | -65.39 | 3.79 | 0.06 | 0.04 | 22.72 | -92.00 |
| 4 | W | 100.00 | 167.53 | -67.53 | 1.09 | 0.06 | 0.03 | 25.24 | -93.95 |

## References

1. Ali, A. Biodiversity–ecosystem functioning research: Brief history, major trends and perspectives. Biol. Conserv. 285, 110210 (2023).

2. Cardinale, B. J. et al. Biodiversity loss and its impact on humanity. Nature 486, 59–67 (2012).

3. Tilman, D., Isbell, F. & Cowles, J. M. Biodiversity and Ecosystem Functioning. Annu. Rev. Ecol. Evol. Syst. 45, 471–493 (2014).

4. van der Plas, F. Biodiversity and ecosystem functioning in naturally assembled communities. Biol. Rev. 94, 1220–1245 (2019).

5. Hector, A. et al. Plant diversity and productivity experiments in European grasslands. Science 286, 1123–1127 (1999).

6. Tilman, D., Wedin, D. & Knops, J. Productivity and sustainability influenced by biodiversity in grassland ecosystems. Nature 379, 718–720 (1996).

7. Mayor, S. et al. Diversity–functioning relationships across hierarchies of biological organization. Oikos 2024, 1–13 (2024).

8. Pickett, S. T. A. & Cadenasso, M. L. Landscape ecology: spatial heterogeneity in ecological systems. Science 269, 331–334 (1995).

9. Wiens, J. A. Landscape mosaics and ecological theory. in Mosaic Landscapes and Ecological Processes (eds. Hansson, L., Fahrig, L. & Merriam, G.) 1–26 (Springer, Dordrecht, 1995).

10. Zonneveld, I. S. The land unit — a fundamental concept in landscape ecology, and its applications. Landsc. Ecol. 3, 67–86 (1989).

11. Peters, D. P. C. & Goslee, S. C. Landscape diversity. in Encyclopedia of Biodiversity (ed. Levin, S. A.) 645–658 (Elsevier, New York, 2001).

12. Duffy, J. E. Why biodiversity is important to the functioning of real-world ecosystems. Front. Ecol. Environ. 7, 437–444 (2009).

13. Jochum, M. et al. The results of biodiversity–ecosystem functioning experiments are realistic. *Nat*. Ecol. Evol. 4, 1485–1494 (2020).

14. Alsterberg, C. et al. Habitat diversity and ecosystem multifunctionality—the importance of direct and indirect effects. Sci. Adv. 3, (2017).

15. Mayor, S. et al. Landscape diversity promotes landscape functioning in North America. Commun. Earth Environ. 6, 1–9 (2025).

16. Oehri, J., Schmid, B., Schaepman-Strub, G. & Niklaus, P. A. Terrestrial land-cover type richness is positively linked to landscape-level functioning. Nat. Commun. 11, 1–10 (2020).

17. Commission for Environmental Cooperation (CEC). 2015 Land Cover of North America at 30 Meters. North American Land Change Monitoring System. https://www.cec.org/north-american-environmental-atlas/land-cover-30m-2015-landsat-and-rapideye/ (2020).

18. Schmid, B., Baruffol, M., Wang, Z. & Niklaus, P. A. A guide to analyzing biodiversity experiments. J. Plant Ecol. 10, 91–110 (2017).

19. Metzger, M. J. et al. A high-resolution bioclimate map of the world: a unifying framework for global biodiversity research and monitoring. Glob. Ecol. Biogeogr. 22, 630–638 (2013).

20. Loreau, M. & Hector, A. Partitioning selection and complementarity in biodiversity experiments. Nature 412, 72–76 (2001).

21. Loreau, M., Mouquet, N. & Holt, R. D. Meta-ecosystems: a theoretical framework for a spatial ecosystem ecology. Ecol. Lett. 6, 673–679 (2003).

22. Yachi, S. & Loreau, M. Biodiversity and ecosystem productivity in a fluctuating environment: The insurance hypothesis. Proc. Natl. Acad. Sci. 96, 1463–1468 (1999).

23. Isbell, F. et al. Quantifying effects of biodiversity on ecosystem functioning across times and places. Ecol. Lett. 21, 763–778 (2018).

24. Gutman, G., Skakun, S. & Gitelson, A. Revisiting the use of red and near-infrared reflectances in vegetation studies and numerical climate models. *Sci*. Remote Sens. 4, 100025 (2021).

25. Oehri, J., Schmid, B., Schaepman-Strub, G. & Niklaus, P. A. Biodiversity promotes primary productivity and growing season lengthening at the landscape scale. Proc. Natl. Acad. Sci. 114, 10160–10165 (2017).

26. Isbell, F. et al. Biodiversity increases the resistance of ecosystem productivity to climate extremes. Nature 526, 574–577 (2015).

27. Tilman, D., Reich, P. B. & Knops, J. M. H. Biodiversity and ecosystem stability in a decade-long grassland experiment. Nature 441, 629–632 (2006).

28. Schnabel, F. et al. Species richness stabilizes productivity via asynchrony and drought-tolerance diversity in a large-scale tree biodiversity experiment. Sci. Adv. 7, eabk1643 (2021).

29. Yu, Q. et al. Species dominance rather than species asynchrony determines the temporal stability of productivity in four subtropical forests along 30 years of restoration. For. Ecol. Manag. 457, 117687 (2020).

30. Mishra, A. K. & Singh, V. P. A review of drought concepts. J. Hydrol. 391, 202–216 (2010).

31. Muñoz-Sabater, J. et al. ERA5-Land: a state-of-the-art global reanalysis dataset for land applications. *Earth Syst*. Sci. Data 13, 4349–4383 (2021).

32. Huang, Y. et al. Impacts of species richness on productivity in a large-scale subtropical forest experiment. Science 362, 80–83 (2018).

33. Reich, P. B. et al. Impacts of Biodiversity Loss Escalate Through Time as Redundancy Fades. Science 336, 589–592 (2012).

34. Wagg, C. et al. Biodiversity–stability relationships strengthen over time in a long-term grassland experiment. Nat. Commun. 13, 7752 (2022).

35. Brophy, C. et al. Major shifts in species’ relative abundance in grassland mixtures alongside positive effects of species diversity in yield: a continental-scale experiment. J. Ecol. 105, 1210–1222 (2017).

36. van Ruijven, J. & Berendse, F. Diversity–productivity relationships: Initial effects, long-term patterns, and underlying mechanisms. Proc. Natl. Acad. Sci. 102, 695–700 (2005).

37. Stockdale, C. A., Macdonald, S. E. & Higgs, E. Forest closure and encroachment at the grassland interface: a century-scale analysis using oblique repeat photography. Ecosphere 10, e02774 (2019).

38. Scherer-Lorenzen, M. et al. Pathways for cross-boundary effects of biodiversity on ecosystem functioning. Trends Ecol. Evol. 37, 454–467 (2022).

39. Hanski, I. Metapopulation dynamics. Nature 396, 41–49 (1998).

40. Leibold, M. A. et al. The metacommunity concept: a framework for multi-scale community ecology. Ecol. Lett. 7, 601–613 (2004).

41. Leibold, M. A., Chase, J. M. & Ernest, S. K. M. Community assembly and the functioning of ecosystems: how metacommunity processes alter ecosystems attributes. Ecology 98, 909–919 (2017).

42. Gounand, I., Harvey, E., Little, C. J. & Altermatt, F. Meta-ecosystems 2.0: rooting the theory into the field. Trends Ecol. Evol. 33, 36–46 (2018).

43. Mendes, C. B. & Prevedello, J. A. Does habitat fragmentation affect landscape-level temperatures? A global analysis. Landsc. Ecol. 35, 1743–1756 (2020).

44. Pau, S., Okin, G. S. & Gillespie, T. W. Asynchronous response of tropical forest leaf phenology to seasonal and El Niño-driven drought. PLoS one 5, e11325 (2010).

45. Schwartz, N. B., Budsock, A. M. & Uriarte, M. Fragmentation, forest structure, and topography modulate impacts of drought in a tropical forest landscape. Ecology 100, e02677 (2019).

46. Sturm, J., Santos, M. J., Schmid, B. & Damm, A. Satellite data reveal differential responses of Swiss forests to unprecedented 2018 drought. Glob. Change Biol. 28, 2956–2978 (2022).

47. Winkler, K., Fuchs, R., Rounsevell, M. & Herold, M. Global land use changes are four times greater than previously estimated. Nat. Commun. 12, 2501 (2021).

48. German Aerospace Center (DLR). TanDEM-X - Digital Elevation Model (DEM) - Global 90m. 10.15489/ju28hc7pui09 (2018).

49. Pettorelli, N. et al. Using the satellite-derived NDVI to assess ecological responses to environmental change. Trends Ecol. Evol. 20, 503–510 (2005).

50. Wang, J., Rich, P. M., Price, K. P. & Kettle, W. D. Relations between NDVI and tree productivity in the central Great Plains. Int. J. Remote Sens. 25, 3127–3138 (2004).

51. Gorelick, N. et al. Google Earth Engine: planetary-scale geospatial analysis for everyone. Remote Sens. Environ. 202, 18–27 (2017).

52. Roerink, G. J., Menenti, M. & Verhoef, W. Reconstructing cloudfree NDVI composites using Fourier analysis of time series. Int. J. Remote Sens. 21, 1911–1917 (2000).

53. Loreau, M. et al. Biodiversity as insurance: from concept to measurement and application. Biol. Rev. 96, 2333–2354 (2021).

54. McKee, T. B., Doesken, N. J. & Kleist, J. The relationship of drought frequency and duration to time scales. in Proc. 8th Conference on Applied Climatology 179–184 (Anaheim, USA, 1993).

55. Butler, D. G., Cullis, B. R., Gilmour, A. R., Gogel, B. G. & Thompson, R. ASReml: Fits Linear Mixed Models using REML. VSN International Ltd (2023).

